# Comparative transcriptomics across nematode life cycles reveal gene expression conservation and correlated evolution in adjacent developmental stages

**DOI:** 10.1101/2020.02.07.938399

**Authors:** Min R. Lu, Cheng-Kuo Lai, Ben-Yang Liao, Isheng Jason Tsai

## Abstract

Nematodes are highly abundant animals with diverse habitats and lifestyles. Some are free-living while others parasitize animals or plants, and among the latter, infection abilities change across developmental stages to infect hosts and complete life cycles. Although parasitism has independently arisen multiple times over evolutionary history, common pressures of parasitism—such as adapting to the host environment, evading and subverting the host immune system, and changing environments across life cycles—have led phenotypes and developmental stages among parasites to converge. To determine the relationship between transcriptome evolution and morphological divergences among nematodes, we compared 48 transcriptomes of different developmental stages across eight nematode species. The transcriptomes were clustered broadly into embryo, larva, and adult stages, suggesting that gene expression is conserved to some extent across the entire nematode life cycle. Such patterns were partly accounted for by tissue-specific genes—such as those in oocytes and the hypodermis—being expressed at different proportions. Although nematodes typically have 3-5 larval stages, the transcriptomes for these stages were found to be highly correlated within each species, suggesting high similarity among larval stages across species. For the *Caenorhabditis elegans-C. briggsae* and *Strongyloides stercoralis-S. venezuelensis* comparisons, we found that around 50% of genes were expressed at multiple stages, whereas half of their orthologues were also expressed in multiple but different stages. Such frequent changes in expression have resulted in concerted transcriptome evolution across adjacent stages, thus generating species-specific transcriptomes over the course of nematode evolution. Our study provides a first insight into the evolution of nematode transcriptomes beyond embryonic development.

## Introduction

Nematodes represent the largest animal phylum on earth and display a vast diversity, with 25,000 species described and about 10,000,000 estimated (Poinar 2011). Their extensive morphological diversity is a reflection of their trophic resources, lifestyles, reproductive strategies, and living environments. Of the free-living nematodes, *Caenorhabditis elegans* is the best-studied model organism in molecular and developmental biology. *Caenorhabditis briggsae*—closely related to *C. elegans* and with an almost identical morphology (Grün et al. 2014) —is also widely used in comparative studies on nematode evolution and development. Additionally, parasitism is ubiquitous in nematodes and has independently arisen at least 18 times during the group’s evolutionary trajectory (Blaxter et al. 1998; Blaxter and Koutsovoulos 2015; Zarowiecki and Berriman 2015; Weinstein and Kuris 2016). Another nematode genus of particular interests is *Strongyloides*, the species of which have a unique life cycle in which they alternate between free-living and parasitic generations. Such alterations make *Strongyloides* a unique and attractive model for studying the evolution of parasitism. A previous study showed that parasitism-associated genes that are expanded and specific to parasitic stages are clustered in specific chromosomal regions, suggesting that they contribute to the regulatory mechanisms of parasite development (Hunt et al. 2016). Some of these parasitism genes expanded across different clades of parasites, indicating convergent evolution at the genomic level (Coghlan et al. 2019); however, the evolutionary relationships among transcriptomes at different stages and parasitic or free-living nematodes remain to be elucidated.

Evolutionary changes occur frequently in organisms through the co-opting of existing traits for new purposes. Which co-opted features change with the emergence of species-specific stages and how they do so at the genetic and regulatory levels are essential questions in evolutionary developmental biology. One theory, the developmental constraint concept, argues that these features limit phenotypic variability and the composition or dynamics of the developmental system (Smith et al. 1985). In nematodes and arthropods, the morphologies and transcriptomes were conserved during mid-embryogenesis between species within the same phylum (Kalinka et al. 2010; Levin et al. 2012). These observations coalesced into the hourglass model. Since evolution and development are two intertwined processes, constraints and variations in a species’ development may have significant impacts on that species’ evolutionary trajectory. A recent study suggested that some gene expressions changed simultaneously across multiple tissues after speciation, leading to correlated patterns of gene expression evolution and causing the genes to group by species in hierarchical clustering (Liang et al. 2018). Studying constraints in the transcriptomes of stages beyond nematode embryogenesis is of tremendous interest, but remains challenging as developments in each stage can be vastly different across intraspecific generations and interspecific morphologies. One of the first experiments comparing transcriptomes of nematode developmental stages beyond embryogenesis was performed by comparing synchronized transitions from embryo to adult stages in *C. elegans* and *C. briggsae* (Grün et al. 2014). It measured fluctuating mRNA and protein expressions across the life stages, and showed that transcript fold changes were conserved during embryo-to-larva transitions. Over the past few years, insights gained from transcriptomic comparisons between developmental stages during and before infection have increased our understanding of parasitism. The recent availability of transcriptomic data from parasitic nematodes (Choi et al. 2011; Stoltzfus et al. 2012; Laing et al. 2013; Hunt et al. 2018; Tanaka et al. 2019) is an exciting resource for identifying the evolution of gene expression throughout development.

The life cycle of nematodes usually consists of one embryo, four to five larval, and one adult stages, which are separated by moulting (Lee 2002; Sommer and Streit 2011). The body size of larvae increases after every moult, eventually reaching sexually mature adult size. Several nematodes have evolved specialised developmental stages, such as a dauer stage whereby the larva undergoes developmental arrest to survive unsuitable conditions, such as a food shortage or high population density. The occurrences of dauer and diapause stages have been associated with gene expression changes in several invertebrates (Flannagan et al. 1998; Bao and Xu 2011; Hand et al. 2016). Unique developmental stages with morphological traits specialized for parasitism also frequently occurred during the evolution of nematode species. These include the sedentary and swollen females in plant-parasitic nematodes and the ensheathed larvae in animal-parasitic nematodes (Lee 2002). In addition, the microfilariae of *Brugia malayi*, which have a morphology very different from any general life stage of nematodes, migrate to and develop in the mosquito, making *Brugia malayi* the intermediate host and transitional insect vector. The transcriptomes of these specialised developmental stages are often distinct to those of previous stages, and these differences in expression mainly come from members of expanded gene families that arose from lineage-specific duplications (Stoltzfus et al. 2012; Baskaran et al. 2015; Hunt et al. 2018).

We hypothesize that, although there is vast phenotypic diversity across the life cycles of different nematodes, these life cycles can be compared and therefore the levels of conservation between gene expression across developmental stages can be quantified. Hence, high throughput sequencing data across nematodes that previously focused on species-specific differences may be further utilized to reveal the conservation of transcriptomes associated with the life cycles. In this study, we compared the transcriptomes of different developmental stages across several nematode species and profiled the conservation of gene expression in whole worms, particular tissues, and biological processes. To investigate if there was conservation at each developmental stage, we estimated the similarities between transcriptomes across developmental stages and clustered based on their similarities. We further quantified these similarities to assess whether specialisation occurred at that developmental stage, and categorised genes into different expression profiles. The frequent changes in orthologue switching profiles between species were revealed and discussed. This study provides the first investigation of developmental conservation across the evolutionary trajectory of multiple nematode species beyond the embryonic stage.

## Results

### Data collection and clustering among intraspecies transcriptomes

We collected independently published transcriptome datasets from five to seven developmental stages in eight nematodes: *C. elegans* and *C. briggsae* (Grün et al. 2014), *Pristionchus pacificus* (Baskaran et al. 2015), *S. stercoralis* (Stoltzfus et al. 2012), *S. venezuelensis* (Hunt et al. 2018), *Haemonchus contortus* (Laing et al. 2013), *B. malayi* (Choi et al. 2011), and *Bursaphelenchus xylophilus* (Tanaka et al. 2019). A summary of the data is shown on Figure 1. Grün et al. (2014) profiled the transcriptomes of two *Caenorhabditis* species during development under the same conditions. The combined dataset consisted of 13 stages with one to six biological replicates per stage and 1.5 to 75 million reads per sample. On average, 82.4% of reads per sample were aligned to corresponding nematode genomes using HISAT2 (Kim et al. 2015) under the same parameters (Supplementary Table S1).

**FIG. 1.**
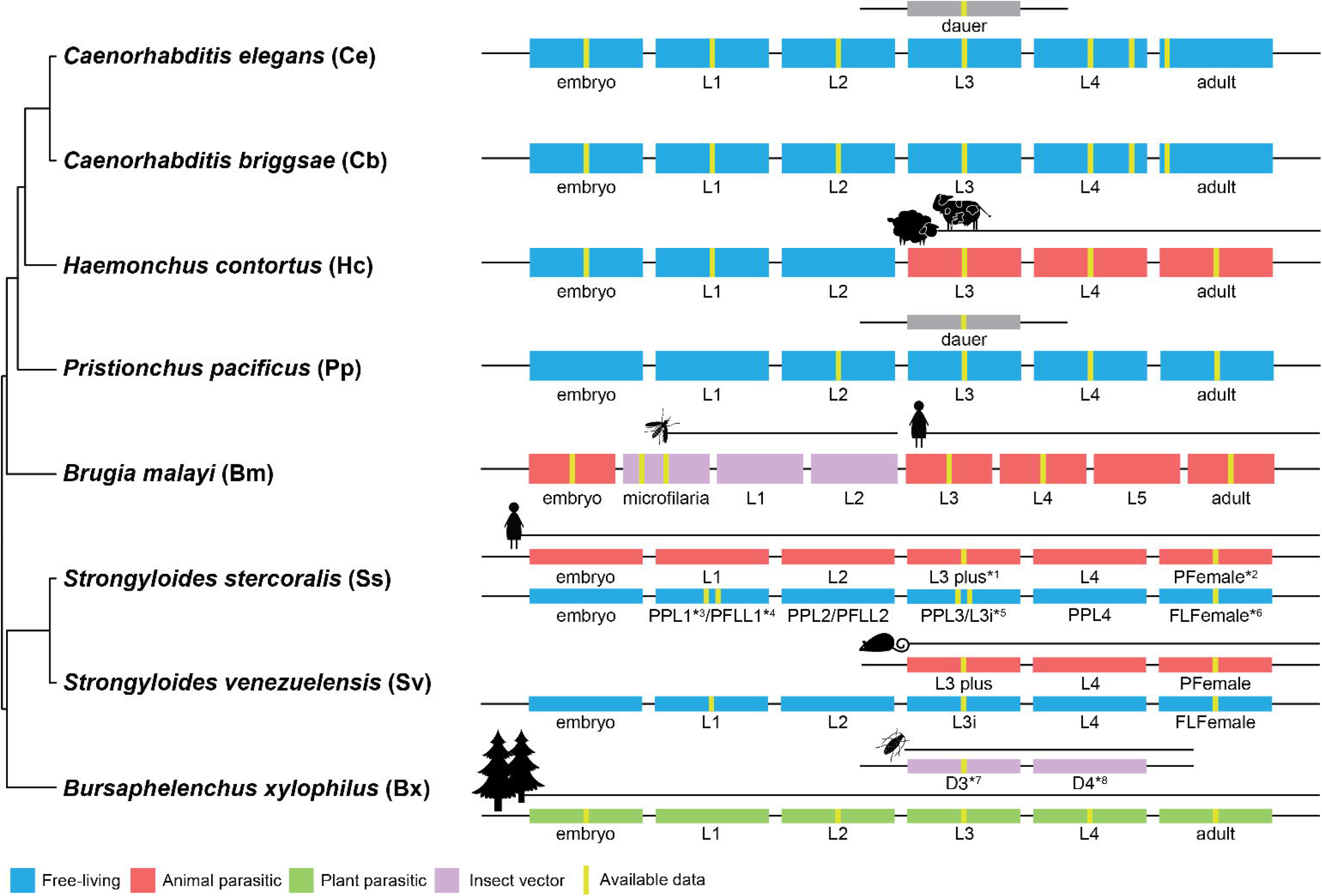
Nematode phylogeny and life cycles. To the left are the phylogenetic relationships among eight nematodes. The abbreviation for each species is shown after the species name, and is used throughout the study. To the right are the life cycles for each species. Different colours correspond to the different nematode lifestyles. The available transcriptome data used in this study are denoted with yellow stripes. Nematode host types are labelled above their life cycles. In total, three free-living, four animal parasitic, and one plant parasitic nematodes were included in this study. (*^1^ L3 plus: Infectious third-stage larva (isolated from host), *^2^ PFemale: Parasitic female, *^3^ PP: Post parasitic, *^4^ PFL: Post free-living, *^5^ L3i: Infectious third-stage larva (isolated from environment), *^6^ FLFemale: Free-living female, *^7^ D3: Dispersal third-stage juvenile (D3), *^8^ D4: Dispersal forth-stage juvenile).

Rather than re-compiling a list of differentially expressed genes between adjacent development stages as done in previous studies (Choi et al. 2011; Stoltzfus et al. 2012; Laing et al. 2013; Grün et al. 2014; Baskaran et al. 2015; Hunt et al. 2018; Tanaka et al. 2019), we were primarily interested in the genome-scale similarities of mRNA expression between development stages. Correlation coefficients between different developmental stages within each species were computed and hierarchically clustered. In general, the transcriptome of the embryo was the most distinct, while those of larval stages were similar to those of their adjacent stages (FIG. 2). In the case of *C. elegans*, each clustered stage—embryo, adult, and larval—were further clustered into early (L1 and L2) and late stages (L3, L4 and LL4) (FIG. 2A). Clustering patterns of larval stages in *P. pacificus* (FIG. 2B) and *B. xylophilus* (FIG. 2C) also supported the early-late partition. This partitioning pattern was not observed for *Haemonchus contortus* (FIG. 2D) or *C. briggsae* (FIG. 2E), possibly due to intra-species variation in the speed of development, which results in worm cultures consisting of individuals with different development stages (Baskaran et al. 2015; Perez et al. 2017). Consistent with previous findings (Baskaran et al. 2015), we also found that specialised phenotypic stages, such as dauer (FIG. 2B), were assigned to its adjacent exit stage (larval L4). In addition, the larval stages were divided into two branches: presence and absence of infectious ability in *S. stercoralis* (FIG. 2F). Similar patterns were found in parasitic larval stages of *S. venezuelensis* (FIG. 2G) and the insect transition stages of *B. malayi* (FIG. 2H). These data suggested that factors associated with parasitism are not as important as factors associated developmental stages in determining similarities of transcriptomes that were analysed. To compare transcriptomes among stages and different species, orthology between genes was first assigned into a total of 15,835 orthogroups, including 2,548-6,736 single copy orthologues across 28 species-pair comparisons.

**FIG 2.**
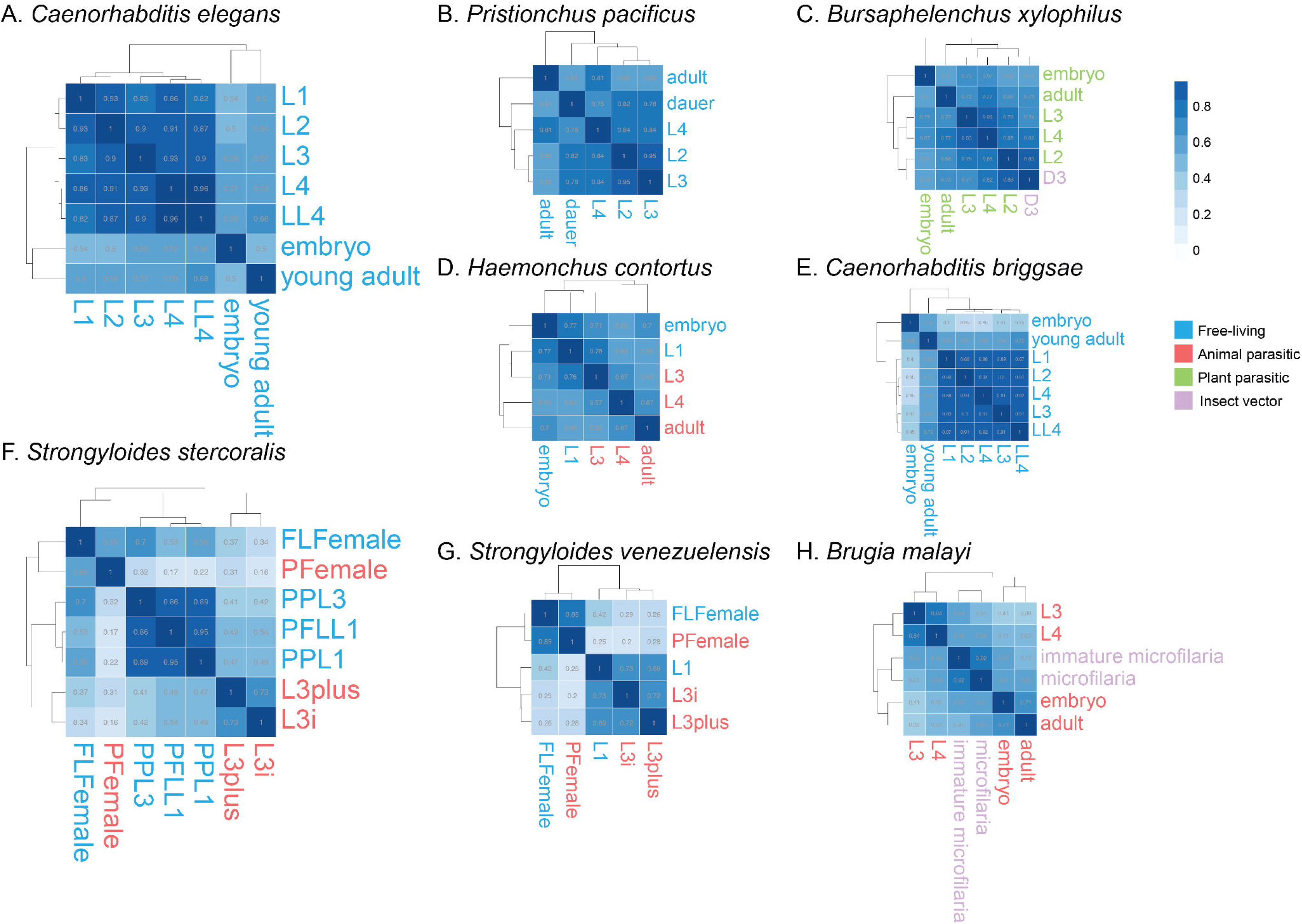
Transcriptome correlation among nematode developmental stages. Results of hierarchical clustering of Pearson correlations at different stages in single species. The number of genes included in the analysis for each species are as follows: *C. elegans* 15,156; *C. briggsae* 16,606; *P. pacificus* 19,137; *S. stercoralis* 9,823; *S. venezuelensis* 12,678; *B. xylophilus* 12,259; *B. malayi* 8,899; *H. contortus* 14,572.

### Transcriptomes across nematode development were clustered into embryo, larval, and adult stages

We first computed the correlation between *C. elegans*-*C. briggsae* single copy orthologues across development stages. The development stages were clustered into embryo, larva, and adult stages (FIG. 3A). Interestingly, expressions across five larval stages were grouped by species, i.e., all *C. elegans* larval stages were grouped together before clustering with the *C. briggsae* larval stages. As this particular dataset was performed under synchronized conditions (Grün et al. 2014), we included additional published data from modEncode (Celniker et al. 2009), which contains transcriptomes from multiple developmental stages (Supplementary Table S3). The same clustering was even observed (Supplementary FIG. S1), even when gene families were also included (Supplementary FIG. S2), demonstrating that such clustering was robust beyond potential batch effects from different studies.

**FIG. 3.**
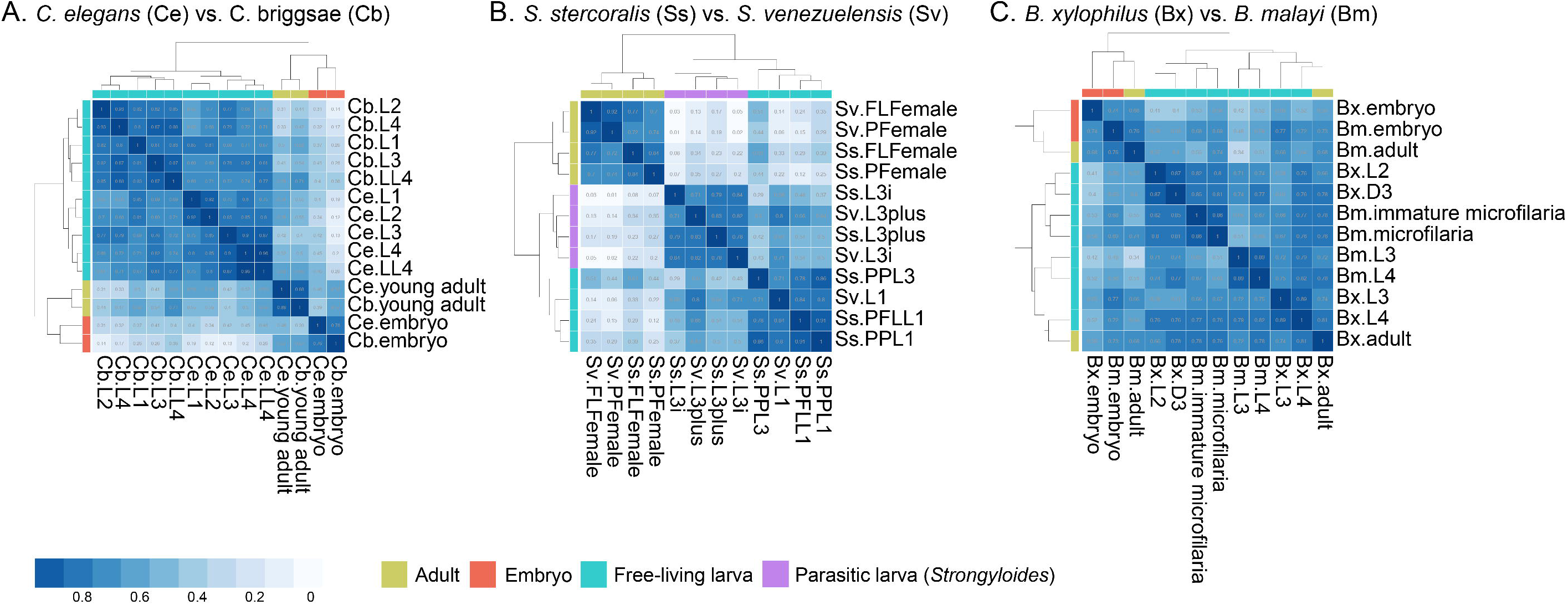
Correlation clustering across nematodes. Hierarchical clustering of Pearson correlation of different stages between (A.) two *Caenorhabditis* species, (B.) two *Strongyloides* species and (C) *B. xylophilus* and *B. malayi*. 6,736, 5,394 and 3,109 one to one orthologues were included in the analysis, respectively. Parasitic larva of *Strongyloides* and the other larva stages were labelled separately.

Similar clustering patterns were also observed in the soil-transmitted gastrointestinal parasitic nematode *Strongyloides* (FIG. 3B). These nematodes are particularly interesting because they alternate between free-living and parasitic generations (Hunt et al. 2016; Hunt et al. 2018). The transcriptomes of *S. stercoralis* (which parasitizes humans) and *S. venezuelensis* (which parasitizes rats) were divided into larva and adult stages, but differences were observed in each cluster. Free-living and parasitic female adult stages of each species were grouped together before clustering into a major ‘adult’ group. Transcriptome clusters in the larval branch were instead separated by lifestyles, consistent with the observation that infection ability was already available in their common ancestor (Hunt et al. 2016) and was conserved despite having different hosts.

We applied the same analysis to all species pairs to systematically compare gene expressions across nematodes (Supplementary FIG. S3). Of the 28 possible combinations, clustering of developmental stages in 13 species pairs again revealed three groups corresponding to three broad developmental processes from embryo to adulthood. We found a lower number of orthologue pairs inferred from species of different genera (2,548-4,295) than pairs from the same genus (5,394-6,736), and such comparisons partitioned the larval stages further into early and late phases. This was the case for the *C. elegans*-*B. xylophilus* and *C. elegans*-*H. contortus* comparisons, but not *C. elegans*-*C. briggsae*. We were concerned that different pairs of orthologue sets would influence the analysis results, and therefore repeated the *C. elegans* and *C. briggsae* clustering with different one-to-one orthologue lists (*C. elegans-B. xylophilus* and *C. elegans-H. contortus*). The clustering results remained identical (Supplementary FIG. S4, FIG. 3A), suggesting transcriptome independence between *C. elegans*-*C. briggsae* larval stages.

Similar environmental pressures lead to recurrent cases of convergence in common genomic adaptations of nematodes of different ancestries (Coghlan et al. 2019). We found one case of transcriptome convergence in the *B. malayi-B. xylophilus* comparison (FIG. 3C). Both species are obligate parasites dispersed by insect vectors. The microfilaria of all stages shared by *B. malayi*-*B. xylophilus* followed the three-theme pattern of embryo, adult and larval stages—except adults of *B. xylphilus*, which were more similar to the late larvae. The late larval stages tended to be grouped by species. In addition, the immature microfilaria and microfilaria congregated with the L2 and D3 larval stages of *B. xylophilus*. First observed by Patrick Manson in the nineteenth century, the microfilaria of *B. malayi* is a unique larval form that allows it to move from the human host’s bloodstream to the intestine of the intermediate vector mosquito. Conversely, *B. xylophilus* is a plant parasitic nematode responsible for pine wilt disease. When faced with an environment high in nematode density and low in food supplies, the *B. xylophilus* L2 larva moults to form an alternative third stage dispersal juvenile (D3), which is taken up by longhorn beetles and transmitted to another healthy host tree (Tanaka et al. 2019). These distinct larval forms adapted to the insect vector were derived independently in their evolution, suggesting transcriptome convergence.

The proportion of different cell types composing the worm body changed dynamically during development. To determine if a similar class of orthologues was also overexpressed in specific tissues throughout development, we dissected the entire worm transcriptome to focus on gene expression in the hypodermis and oocytes. We first obtained 209 and 172 genes involved in these two processes in *C. elegans* based on one previous study of mutants that cause defects in germ cell proliferation (Reinke et al. 2004) and another on cell type-specific RNAseq (Spencer et al. 2011), respectively. We then obtained their orthologues and corresponding expression values in other species. Finally, proportions of the transcriptome from orthologues of *C. elegans* genes in each stage were found to range from 0.2-18% (FIG. 4A-4B). We identified significant differences in transcriptome proportions across developmental stages (embryo, early larva, late larva, and adult; Wilcoxon ranked sum test; FIG. 4C-D). Except for the post parasitic L1 and post free-living L1 stages, expression in the hypodermis was highest during the late larval and embryo stages, followed by the early larval and lowest during the adult stages (FIG. 4A). This is consistent with the finding that mutants with defects in genes involved in hypodermis development produced arrested embryos or larvae (Riddle et al. 1997). Orthologues of *C. elegans* genes that participated in oogenesis had the opposite expression trend, and were expressed the least in the larval stages (FIG. 4C). Interestingly, the immature and mature microfilaria are distinct larval forms in *B. malayi* and also had high expression proportions, which makes sense as this stage comes immediately after the embryo stage and the sheath originates in the envelope of the embryo.

**FIG. 4.**
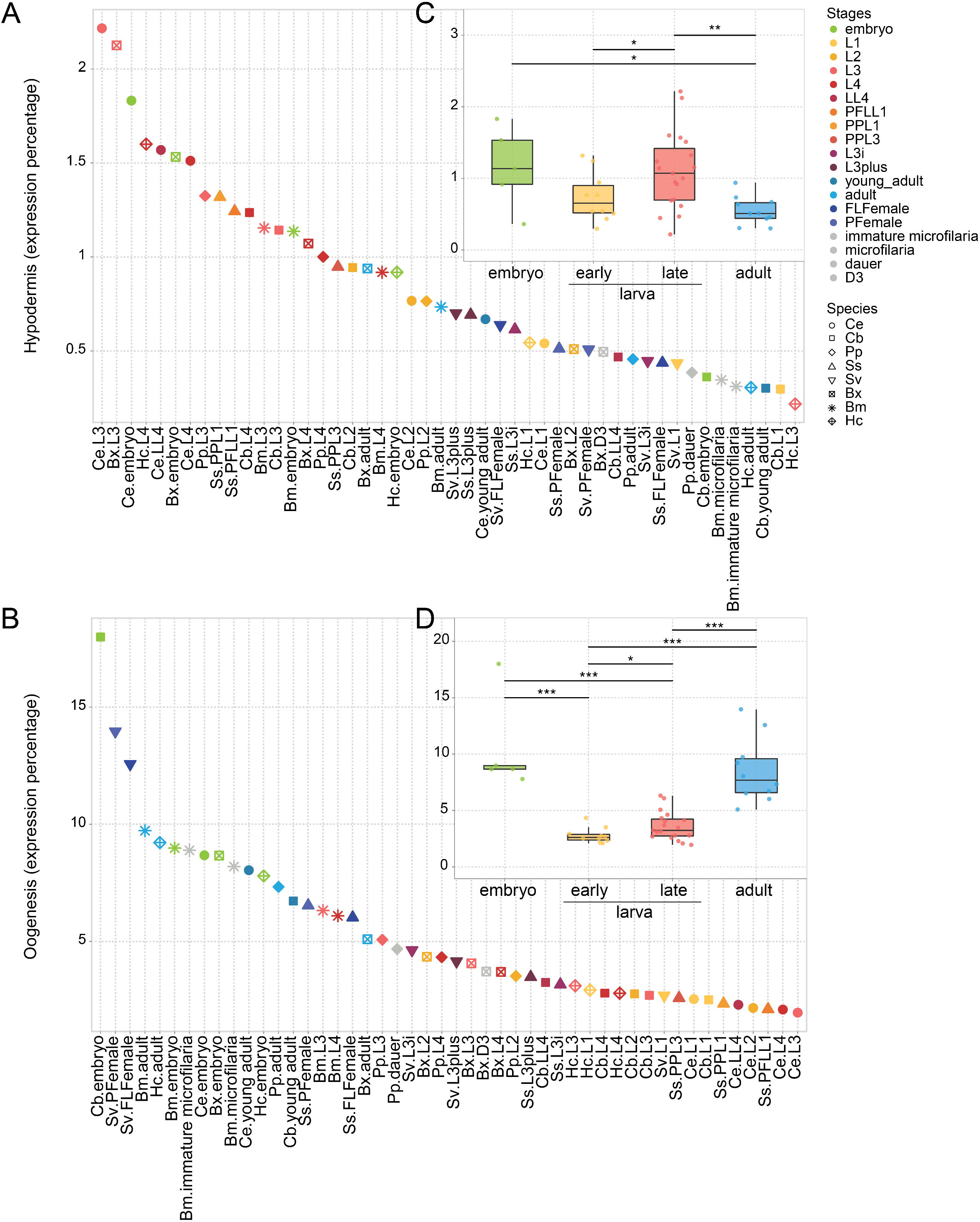
Gene expression in specific gene sets. Proportion of gene expression specifically expressed in (A) hypodermis or (B) during oogenesis compared to the rest of transcriptome. The relative expression level on the y-axis was calculated from the proportion of subset gene expressions in all of the one-to-one orthologues’ expression. The stages across nematodes were assigned to five developmental categories: embryo (green), early larva (yellow), late larva (red), adult (blue), and other(grey). Stages in the ‘other’ group were species-specific stages and excluded in the following analysis. The upper-right figures showed the proportional differences among four different developmental categories in (C) hypodermis or (D) during oogenesis. Wilcoxon rank sum test was performed between each category (p value * < 0.05, ** < 0.01, *** < 0.001).

### Differential levels of correlated transcription evolution (LCE) in nematodes

To further quantify the similarities in the transcriptome across developmental stages, we estimated the level of correlated transcriptome evolution (LCE) (Liang et al. 2018) in stage pairs across species. LCE is a statistical model developed by Liang *et al*. that measures the correlation between transcriptomes by estimating the average correlation across all genes (Liang et al. 2018). High LCE (0.32-0.75) was observed between different larval stages in two *Caenorhabditis* species. In contrast, LCE was 0.01-0.27 for the embryo stage, which overlaps with the LCE bound of 0.235 retrieved from the simulated stage independence (Liang et al. 2018; FIG. 5A). These observations were consistent with the aforementioned finding that different larval stages clustered by species, suggesting that gene expressions in larval stages are correlated. In addition, the finding that LCE was lowest between the embryo and any other stage indicated that the gene expression during embryonic development was distinct from those during larval development and adulthood. In *Strongyloides*, the LCE values for the six comparisons between larva and adult stages were lower than those for all the other comparisons. LCE values in the comparison between parasitic and free-living females, and the infectious L3 and L3+ (L3 collected from host), were 0.64 and 0.36, respectively (FIG. 5B). Consistent with the clustering analysis, LCE results suggested that different lifestyles in adults seemed to have evolved concertedly and not individually.

**FIG. 5.**
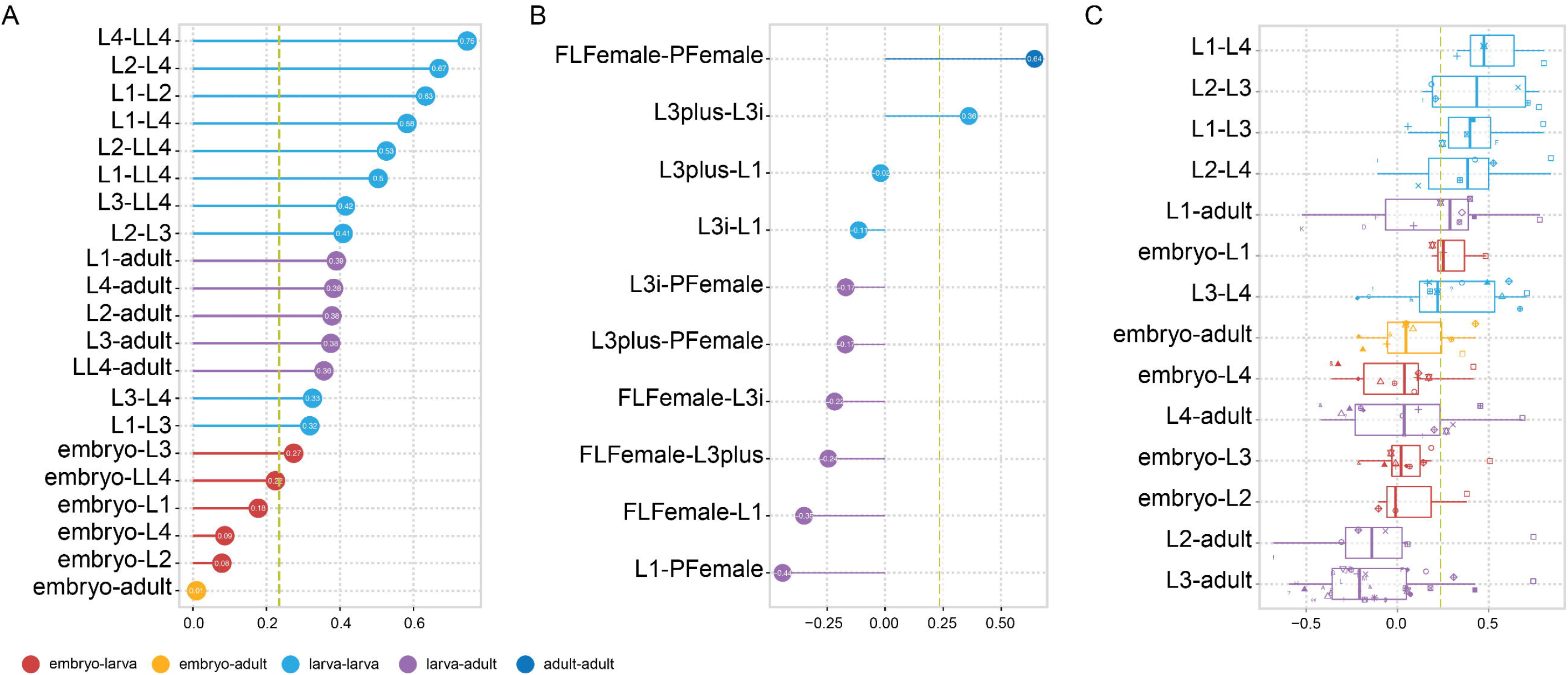
Estimates of levels of correlated evolution (LCE) (A.) LCE between *C. elegans* and *C. briggsae* in 7 developmental stages. (B.) LCE between *S. stercoralis* and *S. venezuelensis*. Free-living L3 (PPL3) and L1 (PFLL1) in *S. stercoralis* were excluded. The PPL1 stage in *S. stercoralis* was assigned as the L1 stage in the comparison because it has the same post parasitic features as the L1 stage in *S. venezuelensis*. (C.) LCE among all the species. The parasitic and free-living generations in *Strongyloides* were separated to compare them to developmental stages in other species. Stage comparisons with fewer than three species pairs were excluded. The lowest LCE value with theoretical p-value < 0.05 was 0.235. Colours indicate the different categories of stage comparison. All larval stages were classified into larva categories to test the level of co-evolution.

Next, we sought to determine whether concerted evolution occurred across nematodes of different genera (FIG. 5C). Species-specific developmental stages, such as microfilaria in *B. malayi* and D3 in *B. xylophilus*, were excluded because they lacked equivalent stages in the other species. Four of the five comparisons between larval stages showed high LCEs (0.27 to 0.53) whereas six of the eight comparisons between larva and embryo or adult had low LCEs (−0.15 to −0.11). Co-evolution of larval stages within a species seemed to be a general phenomenon in nematodes. Of the larval stages, L3 had on average the lowest LCE, while the other stages have average LCEs of 0.38 to 0.51. This suggested that L3 was the most individualized stage. In parasitic nematodes, infectious ability was frequently observed in the third larval stage. This preference for gaining parasitic capabilities provoked the question of whether L3 in parasitic nematodes has evolved independently since early evolutionary history. Our findings on the LCE of parasites suggested that the answer is no, as L3 was not individualized and instead evolved in concert with the adjacent stages after L3 (Supplementary FIG. S5).

### Orthologues are frequently expressed in different developmental stages

To further quantify the differences in transcriptomes among species, we ranked the genes based on relative expression using the Z-score angle-sorted value index for temporal sorting (ZAVIT) (Levin et al. 2016). Next, each gene in each species was categorized based on its ranking and differential expression analysis across developmental stages (FIG. 6A, Supplementary Table S4-5). The expression profiles for *C. elegans* and *C. briggsae* were similar, both showing a stage-like pattern in the embryo, all larval, and adult stages (Supplementary FIG. S6A). The majority of genes were expressed in multiple stages (72.5% and 63.4% in *C. elegans* and *C. briggsae*, respectively). To examine whether there is an evolutionary preference for any particular category of genes, we assessed the differences in expression categories between species. Interestingly, only just half (50.03%) of *C. elegans* orthologues were expressed in the same category in *C. briggsae* (FIG. 6A); the proportion of genes that shifted across developmental profiles varied from 20.6% (in embryo + larva) to 86.1% (in larval + adult). Interestingly, genes of the orthologue pairs expressed in the latter category in both *Caenorhabditis* species exhibited ratios of the nonsynonymous substitution rate to the synonymous substitution rate (*d*_N_/*d*_S_)significantly lower than four other expression profiles (FIG. 6C), implying stronger purifying selection on coding sequences of genes that maintained the same role, despite being in the category with the highest switching. Enrichment of Gene Ontology revealed the significant terms including the “small molecule metabolic process” and “purine nucleoside monophosphate metabolism” (Supplementary Table S6). Small-molecular signalling has been extensively studied in *C. elegans* for its important roles across multiple aspects of development and behaviour (Ludewig 2013) (ref, while purine homeostasis was recently revealed to be necessary for developmental timing in *C. elegans* (Marsac et al. 2019). Both conservation of expression category and higher purifying selection of these genes further imply their functionally importance across *Caenorhabditis* genera.

**FIG. 6.**
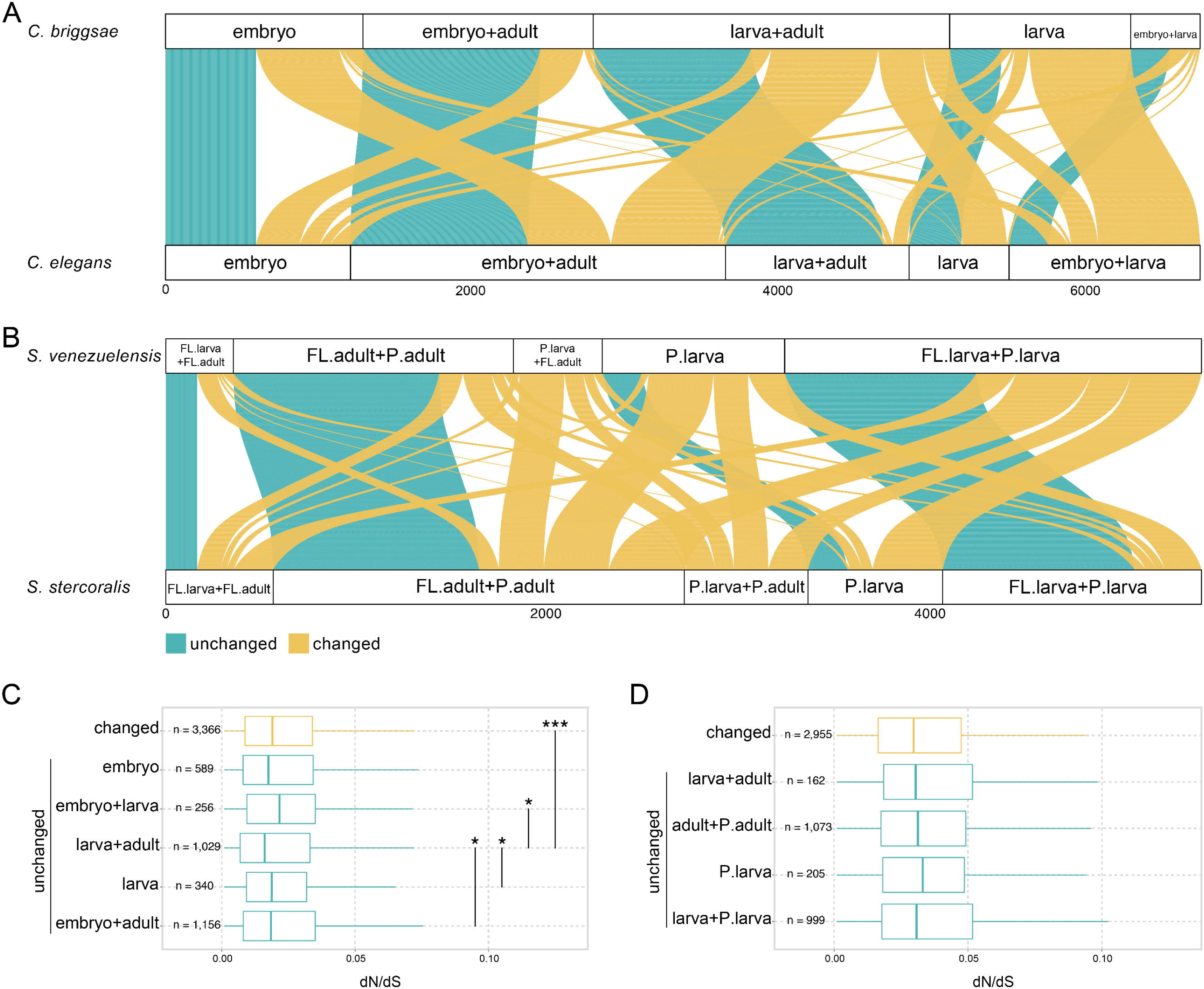
Expression profiles and sequence divergences during development in species of *Caenorhabditis* and *Strongyloides*. Expression profiles of (A.) *C. elegans* and *C. briggsae;* and (B) *S. stercoralis* and *S. venezuelensis*. Expression profiles were categorized into five developmental categories in each species. Sequence divergence of different expression categories in (C.) *C. elegans* and *C. briggsae;* (D.) *S. stercoralis* and *S. venezuelensis*. The five boxes underneath include the genes with the same expression profiles in both species. Wilcoxon test was performed to test the difference in dN/dS between categories. Asterisks were used to represent the p-value (* < 0.05, ** < 0.01, *** < 0.001). Comparisons with p-value > 0.05 were not labelled.

Expressions patterns in free-living and parasitic life cycles were complex and included multiple possible combinations, so we first empirically assigned 88.0% and 91.4% of genes in *S. stercoralis* and *S. venezuelensis*, respectively, into four categories based on when they were expressed (FIG. 6B, Supplementary FIG. S6B): in free-living generations, throughout adulthood, parasitic larval stage only, and throughout adult stage. Interestingly, the rest of genes were assigned to expressed in parasitic larval + free living adult or parasitic larval + parasitic adult in *S. venezuelensis* and *S. stercoralis*, respectively. Strikingly, the majority of genes (87.0% in *S. stercoralis* and 82.4% in *S. venezuelensis*) were expressed in more than one developmental phase, while over half (54.8%) of *Strongyloides* orthologues were assigned to different developmental categories. Higher number of *Strongyloides*’ genes were expressed at multiple stages than *Caenorhabditis*, but similar levels of developmental switching. The *d*_N_/*d*_S_ ratios were calculated for each category, and none exhibited a significantly higher ratio than genes that exhibited a different expression category (FIG. 6D).

## Discussion

Very little is known about transcriptome conservation between nematodes beyond the embryonic stage. In this study, we compared the developmental transcriptomes of eight nematodes species with similar developmental stages. These species have many morphological and developmental differences and a diversity of living environments, lifespans, and host types. The transcriptomes used in this study came from datasets from multiple sources, but all clustered into three broad stages (embryonic, larval, and adult) across nematodes’ entire life cycles. One major concern was that batch effects would lead to systematic differences between datasets (Fei et al. 2018) that could not be separated from species effects (Leek et al. 2010). Attempts to treat batch effects would also remove biological signals from species. Despite this caveat, the same pattern was observed when we incorporated transcriptomes of the same stage from multiple sources of *C. elegans*, suggesting that the biological signals were robust enough to make up for the technical variations across different studies. Another challenge presented in our study was that the development stages were incomplete in some nematodes, which led to reduced resolution in our analyses. Imperfect synchronization of worm cultures was observed during the staging protocol in *P. pacificus*, and such issue may be applicable to all non-model species. In addition, the definition of developmental stages beyond embryogenesis to adulthood in nematodes was only loosely defined by moulting. Even in the model *Caenorhabditis*, we still have concerns related to perfect synchronization of developmental stages. For instance, the culturing environment may have been more stressful for one species than the other. Further experimental work across all nematode genera is needed to characterize patterns of conservation across their entire life cycles.

Cell-specific information is critical for deciphering how molecular mechanisms control the phenotypes of multicellular animals. Large scale research on gene expression in mammalian organ development suggests that organs become increasingly more distinct and the breadth of gene expression gradually decreases during development (Cardoso-Moreira et al. 2019). This evidence supports the theory of von Baer, that morphological differences between species increase as development advances (Abzhanov 2013). So far, gene expression has been determined in specific cell types of *C. elegans* by dissecting different worm tissues (Spencer et al. 2011) and machine learning-based predictions (Kaletsky et al. 2018). Across nematodes, we revealed that orthologues of oogenesis and hypodermis specific genes in *C. elegeans* also displayed a shared pattern of transcriptome proportions across developmental stages. Identifying tissue-specific genes in other nematodes using data from *C. elegans* alone will inevitably underestimate the proportions of tissue expression contributing to whole-worm transcriptomes. We attempted to minimise such bias by normalising total gene expression to only one-to-one orthologues. Although one-to-one orthology may capitulate a subset of developmentally conserved genes, we have shown that they provided initial insights into gene expressions during tissue development across nematodes.

We examined the stage clustering further by inferring the level of correlated transcriptome evolution (LCE), a statistical method originally intended to quantify correlation between tissue transcriptomes (Liang et al. 2018). Although the LCE estimation revealed strong concerted evolution between individual larval stages, an alternative explanation may be the imperfect synchronization of worm culture in non-model organisms. For instance, the staging protocol in *P. pacificus* was observed in major developmental transcriptome clustering with a mixture of early larvae, late larvae, and adult (Baskaran et al. 2015). Nevertheless, this may not be the case, at least in the *Caenorhabditis* dataset, as the largest proportion of expression profiles were non-adjacent developmental stages (FIG. 6A). At least half of the genes in nematodes were expressed at multiple stages, and half of their orthologues were found to be expressed at multiple other stages; this suggests that a change in a gene’s expression during development may rapidly lead to transcriptome divergence after speciation.

Using *Strongyloides*, a unique model to investigate the parasitism based on the fascinating features of both free-living and parasitic generation, allowed us to systematically examine the differences between the same developmental stages in different life style strategies. We found that the transcriptomes of *Strongyloides* can be categorized based on developmental stage instead of lifestyle, which is in contrast to the observation that up to 20% of the genes are differentially expressed between parasitic and free-living females (Hunt et al. 2018). The reason behind the two different observations is that that the majority of these differentially expressed genes were duplicated in the *Strongyloides* lineage (Hunt et al. 2018), and we focused on one-to-one orthologues. We have to a certain extent identified a theme of developmental conservation across nematodes, and shown that the specialisation into parasitic stages was the result of duplication events in gene families, as is evident in many nematode genomes (Hunt et al. 2016; Coghlan et al. 2019). We speculate that altering the expression of a gene to adapt to a new environmental niche may take place before genomic innovation without reducing much fitness.

The third larval stage is thought to be a hot spot for obtaining infectious ability (Lee 2002). Gene expressions in the third larval stages were clustered in parasites, but not with all corresponding stages, especially the free-living L3 and diapause ones. This suggests that the similarities in transcriptomes among parasitic stages were not inherited from their common ancestors but through convergent evolution of having similar selection pressures to tolerate the host environment. The results of the expression divergence analysis show that genes tend to be expressed multiple times over the course of the developmental process. We propose that the life cycle of the nematode common ancestor consisted of an embryo stage and an adult stage, with several larval stages in between. The specialised larval stages—such as the dauer, filarial, and sheathed larva stages—may have independently evolved in response to biological requirements over evolutionary time.

Our study adds to recent efforts to sequence and compare genomes across many nematodes (Coghlan et al. 2019) by providing a first step towards revealing life cycle conservation and convergence at the transcriptome level. The most striking pattern was perhaps the finding that some patterns are conserved in species that diverged many millions of years ago and have drastically different lifestyles. Our results also provide initial insights into how ancestral life strategies such as parasitism evolved to become specialised. Future large-scale synchronised experiments across life cycles as well as tissue specific or single cell transcriptomes between nematodes may further elucidate life cycle evolution in nematodes.

## Methods

### RNA-seq mapping and normalization

A description of locations where RNA sequencing (RNA-seq) reads were downloaded is presented in Supplementary Table S1. RNA-seq reads were first trimmed using Trimmomatic (v0.36; parameter: LEADING:5 TRAILING:5 SLIDINGWINDOW:3:15 MINLEN:34; Bolger et al. 2014) to remove the adaptor and leading, tailing, and low quality sequences. Trimmed reads from each species were mapped to a corresponding genome assembly downloaded from Wormbase (ver. WS269; Lee et al., 2018) using HISAT2 (ver. 2.2.1; Kim et al. 2015). Raw gene counts were assigned using featureCounts (v.1.6.3; Liao et al. 2014). The raw counts of orthologous genes in all samples were transformed into TPM (transcripts per million), and the median of replicates were calculated to represent the raw gene expressions of developmental stages in each species. To normalise the data, we initially removed the 25% lowest-expressed genes in each species using the sum of samples. Next, we performed the ‘withinLaneNormalization’ function in EDASeq (v2.18.0) (Risso et al. 2011) to remove GC bias for each gene, and transformed the expressions by log2. Considering that our data were collected from multiple studies, we accounted for study design batch effects using the ‘ComBat’ function from the sva (v3.32.1) (Leek et al. 2016). In the case of species-paired comparisons, both orthologues below the 25% expression category were removed for further analyses. Pearson correlation coefficient of normalised transcriptomes in different developmental stages within and between species were determined using the ‘corr’ function in R (v3.6.0; R Core Development Team 2019). The heat maps of correlation matrices were hierarchically clustered with the average agglomeration method.

### Phylogenetic and evolutionary analysis

Orthology of proteomes from species investigated in this study was inferred using OrthoFinder (v2.2.7; Emms and Kelly 2015). If multiple isoforms exist for a given gene, only the longest or major isoform was chosen for analyses. A maximum likelihood phylogeny was constructed by the concatenated amino acid alignments of 2,205 single copy orthologues across eight nematodes using RAxML (v8.2.11; -m PROTGAMMAILGF -f -a; (Stamatakis 2014) with 500 bootstrap replicates. To calculate sequence-based metrics, sequences of single copy orthologues were retrieved and aligned using TranslatorX (version 1.1; Abascal et al. 2010). We identified the synonymous (dS) and nonsynonymous (dN) substitution rates using Codeml in PAML (v4.9; parameter: runmode=-2, seqtype=1, CodonFreq=3, fix_omega=0; Yang 2007). The 209 oogenesis-enriched genes were defined by (Reinke et al. 2004). The 172 hypodermis-specific genes in *C. elegans* were defined by (Spencer et al. 2011). One-to-one orthologues of these *C. elegans* genes in other species were retrieved. To deal with the differences in orthologue numbers, expression levels of selected genes were normalised to those of one-to-one orthologues with *C. elegans*, generating a relative proportion to represent the expression of specific gene sets.

### Comparative transcriptomic analysis

Levels of correlated evolution (LCE) between transcriptome datasets was calculated according to Liang *et al*., (https://github.com/cloverliang/LCE). We applied the Z-score angle-sorted value index for temporal sorting (ZAVIT) method (Levin et al. 2016) to organize gene expressions across the developmental process. ZAVIT sorted the standardized gene expressions by the relative order of the first two principal components. The standardized expression profiles were obtained by subtracting the mean and dividing by the standard deviation of the orthologous gene expression for all developmental stages. Principal component analysis was performed to standardize gene expression profiles into ordered genes in a circle, and the angle computed by an invert tangent from the origin represented the temporal expression profiles during development. The first gene in the sorted standardized gene expression profile plot was defined according to the sliding window below.

All developmental stages were first categorized into embryo, larva, and adult stages. Additionally, free-living or parasitic stages were considered separated in the *Strongyloides* comparison. This yielded three and four stages in the *Caenorhabditis* and *Strongyloides* comparisons, respectively. Differential expression analyses between developmental stages were performed using DESeq2 (v1.24.0, padj < 0.05; Love et al. 2014). 81.3-82.2% and 63.1-70.2% of genes were first unambiguously sorted into seven and nine expression categories in the *Caenorhabditis* and *Strongyloides* comparisons, respectively, based solely on DESeq2 results. These results were used to further split the ZAVIT-sorted genes into five major expression categories. The boundaries were set by choosing the 5% and 10% quantile of the DEseq2 categories in the *Caenorhabditis* and *Strongyloides* comparisons, respectively. We were interested in the relative order of each gene’s position within the transcriptome, and this approach allowed for the placement of certain genes—e.g., the housekeeping genes, which were expressed throughout development, were sorted relative to the rest of the transcriptome. The starting position of the parasitic larva + parasitic adult expression category in *S. stercoralis* and *S. venezuelensis* were determined solely by the 95% quantile of the adjacent free-living adult + parasitic adult DEseq category, as genes positioned around this region tend to exhibit no significantly different expressions in DEseq2 analyses. Gene Ontology enrichment was performed using topGO (v2.36.0; Alexa and Rahnenfuhrer 2019) with GO annotations downloaded from WormBase.

## Authors contribution

I.J.T conceived the study. M.R.L carried out the majority of analysis with help from C.K.L and B.Y.L. M.R.L and I.J.T wrote the manuscript.

## Competing interests

The authors declare no competing interests.

