## Supplementary Information for "Comparative transcriptomics across nematode life cycles reveal gene expression conservation and correlated evolution in adjacent developmental stages"

**Supplementary Figures**

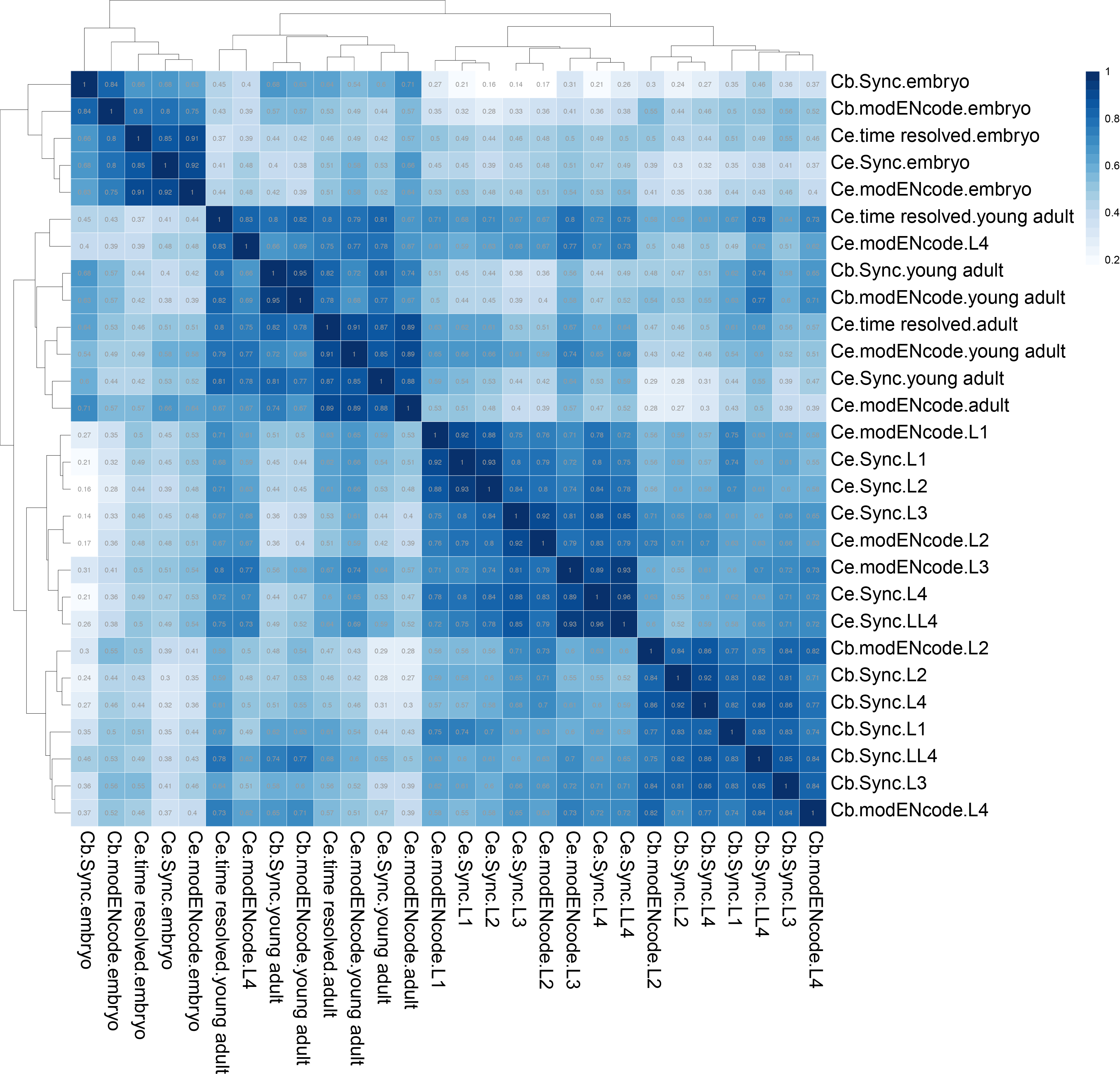

**FIG. S1. Transcriptome clustering of multiple studies between *C. elegans* and *C. briggsae*.**

Despite the dataset used in this study, two independent datasets were added, including one with *C.elegans* and *C. briggsae* data from modENCODE (ref) and the other with high resolution of embryonic and post-embryonic stages in *C. elegans* (ref). These three independent datasets were respectively labeled with Sync, modENCODE and time resolved in the figure. Only the L4 stage of *C. elegans* (modENCODE) did not follow the three theme patterns.

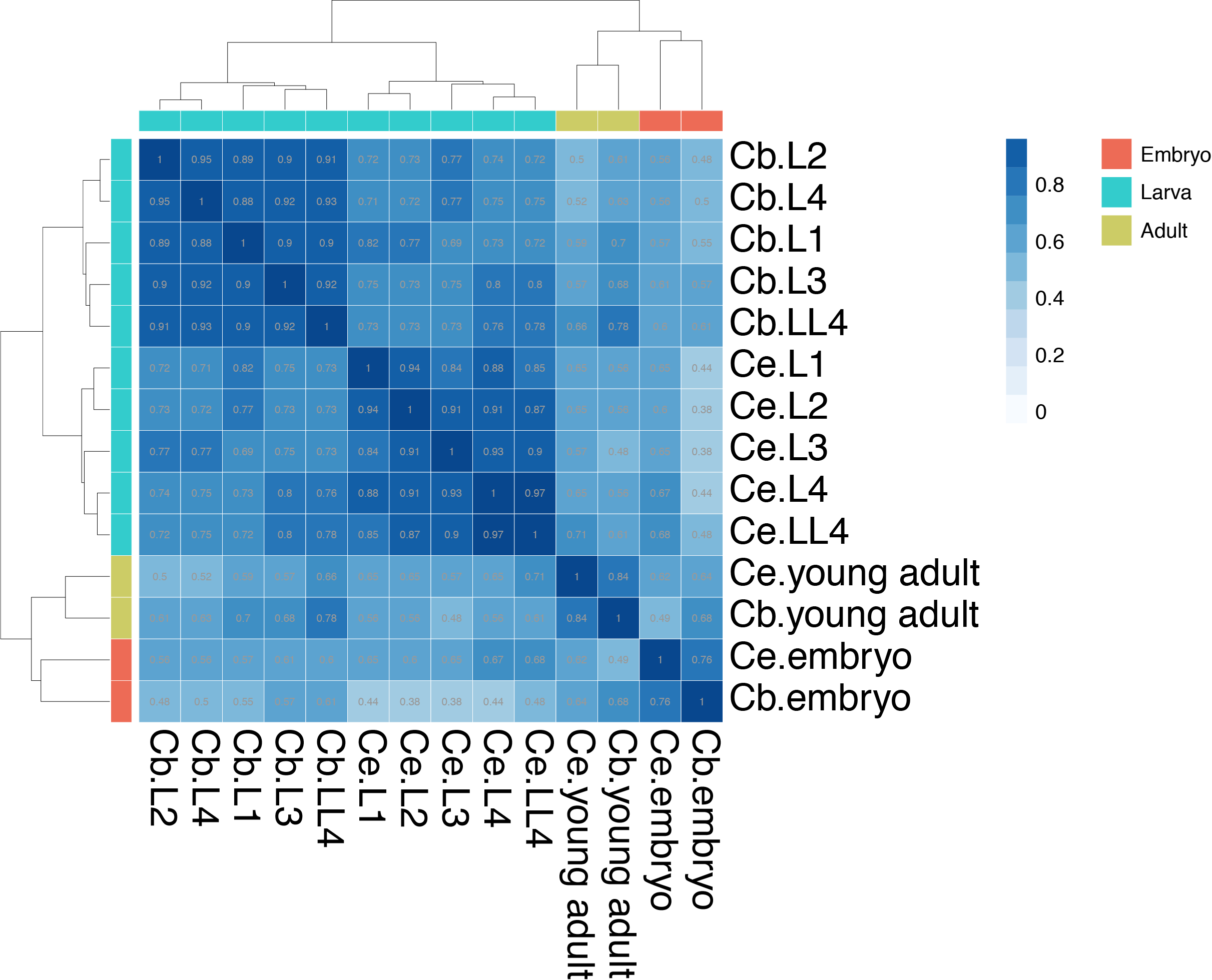

**FIG. S2. Correlation clustering of all orthogroups between *C. elegans* and *C. briggsae*.**

Expressions of gene families, calculated from the summation of TPM in an orthogroup with multiple gene copies. Correlation among 11,512 orthogroups were presented after filtering out 4,323 genes with low expression in *C. elegans* or *C. briggsae*.

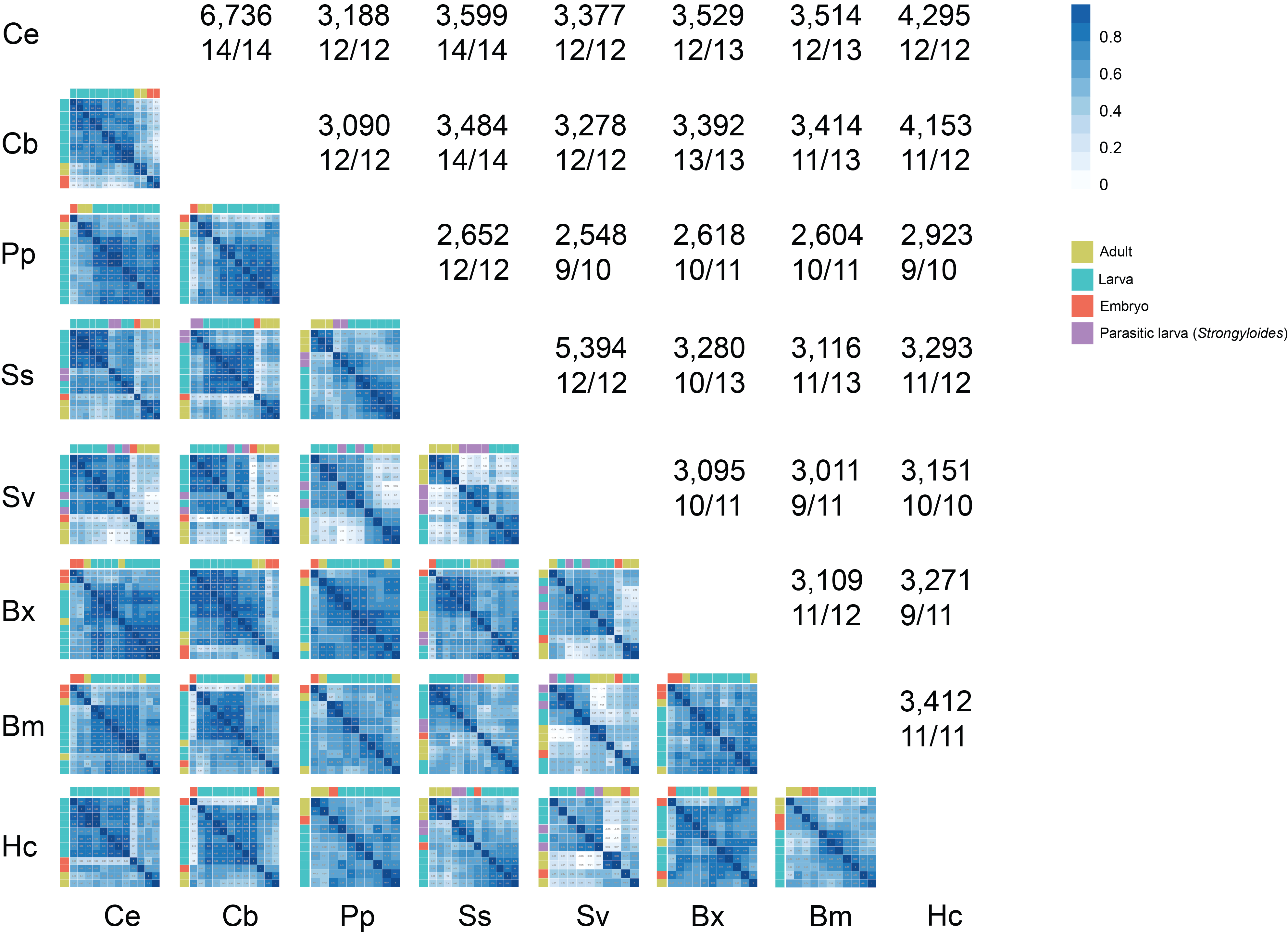

**FIG. S3. Pairwise correlation clustering of nematodes.**

Pairwise comparison of the correlation clustering across eight nematode species were presented. The ratio of stages followed the three themes pattern and the number of the examined one to one orthologues were indicated above each plot. Numbers represent the correlation value of the comparison of the stages in the species pairs.

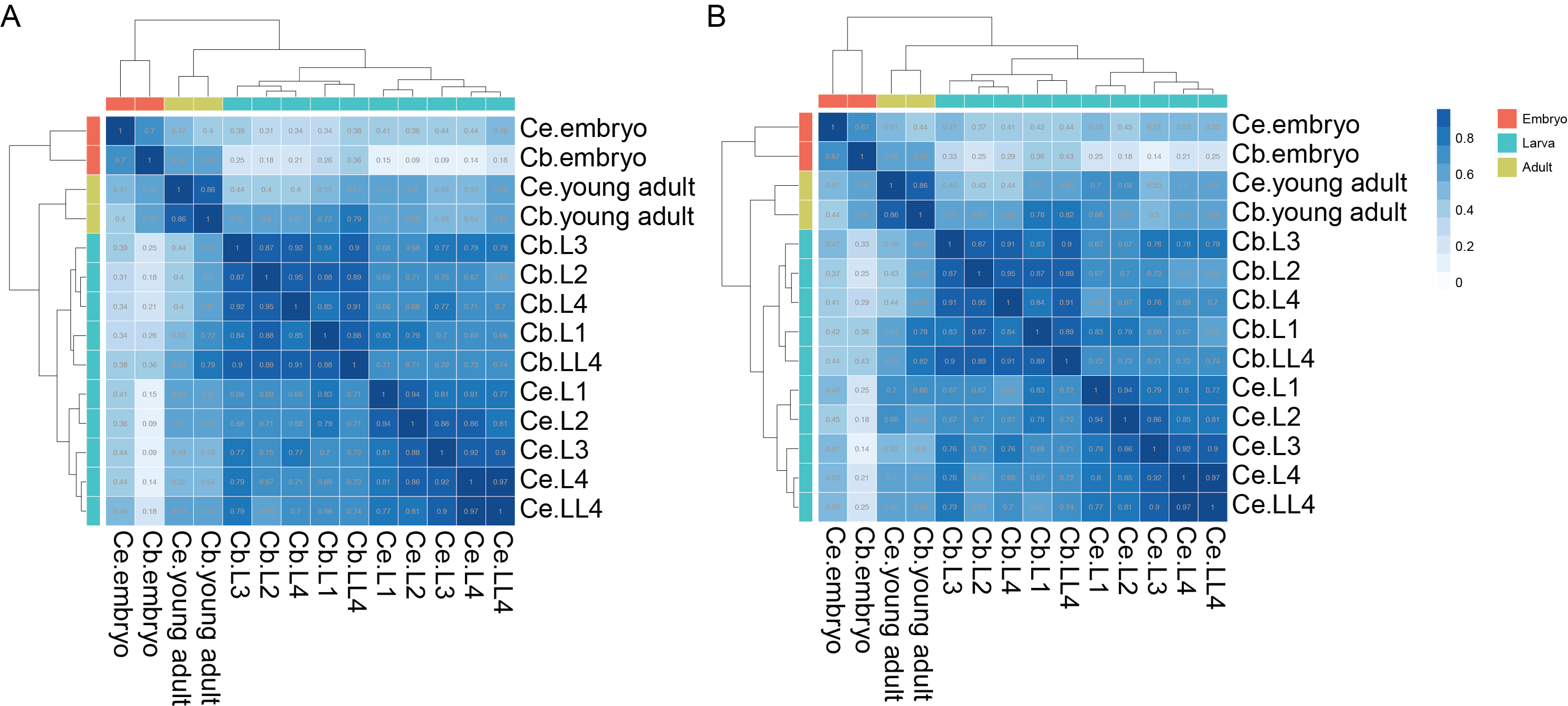

**FIG. S4. Correlation clustering in *Caenorhabditis*.**

(A.) Correlation clustering in *Caenorhabditis* using 4,143 one-to-one orthologue of *C. elegans-H. contortus*. (B.) Correlation clustering in *Caenorhabditis* using 3,393 one-to-one orthologue of *C. elegans*-*B. xylophilus.* Note that some one-to-one orthologues in *C. elegans-H. contortus* and *C. elegans*-*B. xylophilus* where not found in *C. elegans-C. briggsae*.

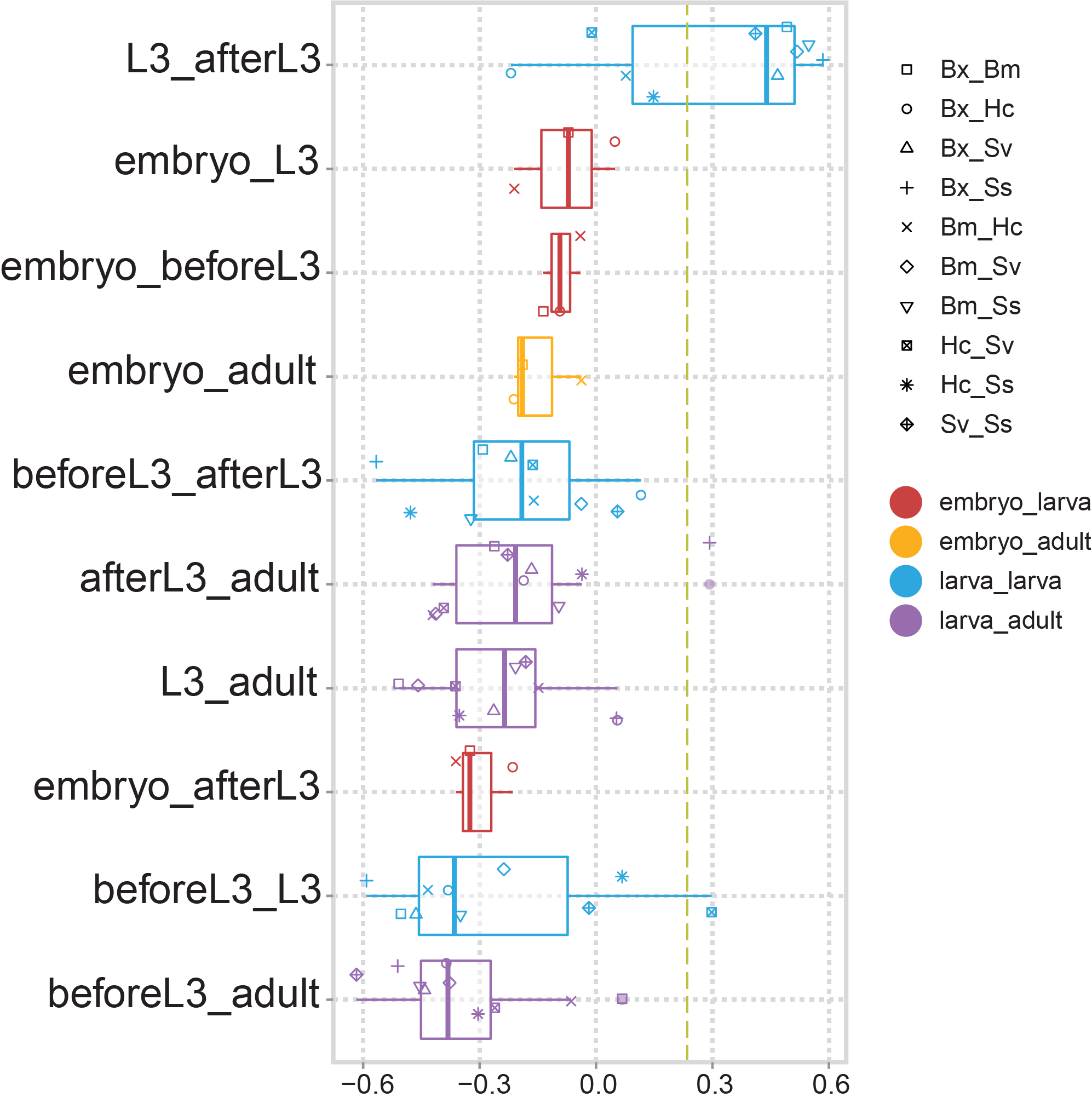

**FIG. S5. LCE estimation of parasitic nematodes.**

2,156 one-to-one orthologues across five species were chosen for LCE estimation in all species comparisons. Five developmental stages were selected in each parasitic nematode species, including embryo, before L3, L3, after L3, and adult. The adjacent stages of infectious L3 were assigned as the before or after L3 stages. Due to the sparsity of our dataset, the before L3 stages were PFLL1 in *S. stercoralis*, L1 in *Strongyloides*, L2 in *B. xylophilus*, microfilariae in *B. malayi*, and L1 in *H. contortus,* while the after L3 stages were L3 plus in *Strongyloides* and L4 in the other three species. In addition, the parasitic females of *Strongyloides* were assigned as adults in this analysis.

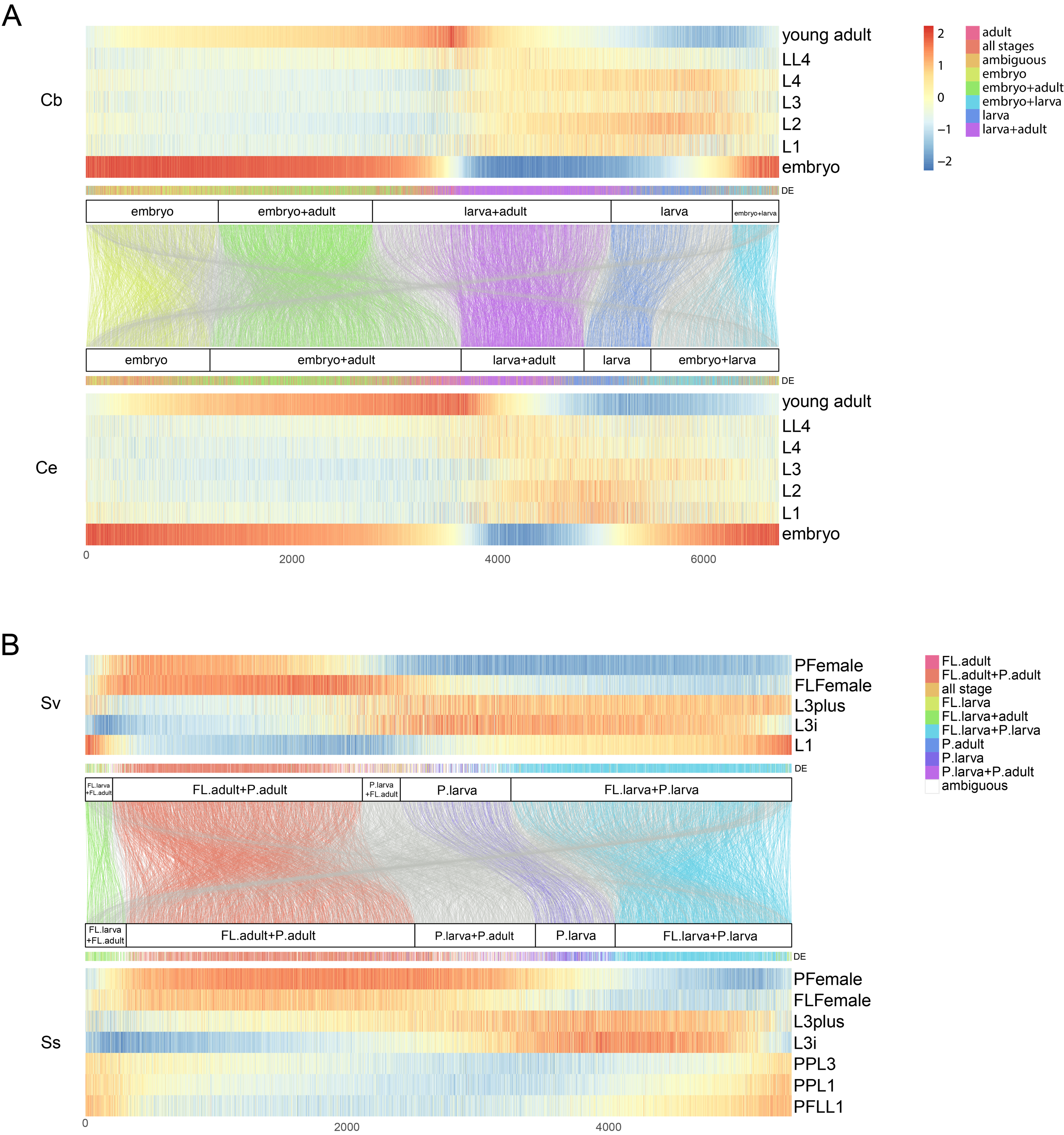

**FIG. S6. Time-sorting expression profile of *Caenorhabditis* and *Strongyloides*.**

Expression profiles were categorized into categories by differential expression and the expression profile in each species. The coloured lines in the middle panel indicated the relative positions of an orthologue between two species. The one-to-one orthologues with different expression profiles in the two species are shown in grey.
